# Statistical and machine learning models for classification of human wear and delivery days in accelerometry data

**DOI:** 10.1101/2020.12.31.424867

**Authors:** Ryan Moore, Kristin R. Archer, Leena Choi

## Abstract

**Purpose:** Accelerometers are increasingly utilized in healthcare research to assess human activity. Accelerometry data are often collected by mailing accelerometers to participants, who wear the accelerometers to collect data on their activity. The devices are then mailed back to the laboratory for analysis. We develop models to classify days in accelerometry data as activity from actual human wear or the delivery process. These models can be used to automate the cleaning of accelerometry datasets that are adulterated with activity from delivery.

**Methods:** For the classification of delivery days in accelerometry data, we developed statistical and machine learning models in a supervised learning context using a large human activity and delivery labeled accelerometry dataset. We extracted several features, which were included to develop random forest, logistic regression, mixed effects regression, and multilayer perceptron models, while convolutional neural network, recurrent neural network, and hybrid convolutional recurrent neural network models were developed without feature extraction. Model performances were assessed using Monte Carlo cross-validation.

**Results:** We found that a hybrid convolutional recurrent neural network performed best in the classification task with an F1 score of 0.960 but simpler models such as logistic regression and random forest also had excellent performance with F1 scores of 0.951 and 0.957, respectively.

**Conclusion:** The models developed in this study can be used to classify days in accelerometry data as either human or delivery activity. An analyst can weigh the larger computational cost and greater performance of the convolutional recurrent neural network against the faster but slightly less powerful random forest or logistic regression. The best performing models for classification of delivery data are publicly available on the open source R package, *PhysicalActivity*.

## Introduction

Recent advances in accelerometers, devices that measure changes in velocity, have led to their increased use in health-care research with applications such as fall detection in the elderly (Bagala et al. 2012), physical activity energy expenditure (Crouter, Churilla & Bassett 2006), and human activity recognition (Kwapisz, Weiss & Moore 2011). The analysis of accelerometry data can present unique challenges due to massive data, participant non-adherence to protocol, and the data often being collected outside of controlled laboratory settings. Additionally, accelerometers are often activated prior to shipment to participants and are not deactivated until they are returned to the laboratory. This process can cause large portions of accelerometry datasets to be recorded while the accelerometers are in transit to the participant or laboratory. An analyst must perform the arduous task of manually removing delivery activity from the dataset. The purpose of this study is to develop models that can accurately classify a given day in an accelerometer dataset as “human wear” or “delivery.” These models can then be used to automate the removal of delivery days in accelerometer datasets before the analysis on human activity.

Prior research has created algorithms for identifying when participants are not wearing the accelerometer (Choi, Liu, Matthews, & Buchowski 2011; Choi, Ward, Schnelle & Buchowski 2012); however, models to classify whether accelerometer activity is due to human wear or motion from delivery have not been developed. Fortunately, the field of human activity recognition is highly applicable to this problem. The primary goal in human activity recognition research is to classify temporal partitions in a dataset in which different activities are performed (Kim, Helal & Cook 2009). Human activity recognition is performed with data from a wide variety of sensors such as video cameras, GPS, heart monitors, and thermometers, and wearable accelerometers are one of the most commonly utilized devices due to recent technical advances in microelectronics and the rich data they provide (Lara & Labrador 2012). The classification of days as human wear or delivery based on differences in acceleration is a similar problem to human activity recognition. Statistical models utilized to classify human activity may be applicable to the development of models to classify accelerometer data as human wear or delivery activity.

In the context of human activity recognition, accelerometer data are traditionally analyzed by first extracting global features such as time between peaks and average acceleration from the temporal intervals of the dataset. By extracting features, massive datasets can be reduced such that regression models or machine learning methods can be used with a reasonable number of variables. Kwapisz et al. (2011) successfully utilized extracted features from accelerometer data to accurately classify 6 human activities with multilayer perceptron (Rosenblatt, 1958) and logistic regression models in the Wireless Sensor Data Mining (WISDM) project. More recently, Ellis et al. (2015) developed a random forest model (Breiman, 2001) that used extracted acceleration features to classify human activity into 4 types with high accuracy. Utilizing feature extraction to develop models is very common in the field of human activity recognition; however, the recent advancement of neural networks and computing power allows direct analysis of the raw data to learn complex features. The convolutional (LeCun et al., 1999) and long short-term memory (LSTM, Hochreiter and Schmidhuber, 1997) neural network architectures have been used in human activity recognition due to their ability to automate the collection of local and temporal features. Ignatov (2018) recently developed a convolutional neural network that learned local spatial features, while simultaneously using extracted global features. Another recent publication (Ordóñez and Roggen, 2016) was successfully able to predict human activity from accelerometer data using a hybrid convolutional LSTM recurrent neural network.

We considered several methods traditionally used in human activity recognition to develop models for the classification of days in an accelerometry dataset as human wear or delivery. Our models were developed in a supervised learning context using a large human activity and delivery labeled accelerometry dataset. We developed logistic regression, mixed-effects logistic regression, random forest, and multilayer perceptron models using extracted features from the dataset. Additionally, we developed convolutional neural network, LSTM recurrent neural network, and hybrid convolutional recurrent neural network models using scaled raw tri-axial accelerometer data as inputs without specific feature extraction. The developed models will be useful for clinicians and researchers interested in classifying accelerometry data as either scientifically relevant human wear days or delivery days to be removed.

## Methods

### Data

The accelerometry dataset used to fit our models is composed of 779 assessments in which 251 participants were mailed a tri-axial Actigraph accelerometer to wear for one week at three time-points during a randomized controlled trial in patients undergoing spine surgery. Actigraph assessments occurred at 6 weeks, 6 months, and 12 months after surgery (Archer, Huag, & Pennings, 2020; Coronado *et al*., 2020). Approximately 54% of the days in the dataset are delivery days, while the remainder are human wear. Physical activity was measured with 1-minute epoch from the x, y, and z axes. An example of accelerometry data from the x-axis over the course of one assessment is shown in **Figure 1**.

**Figure 1:**
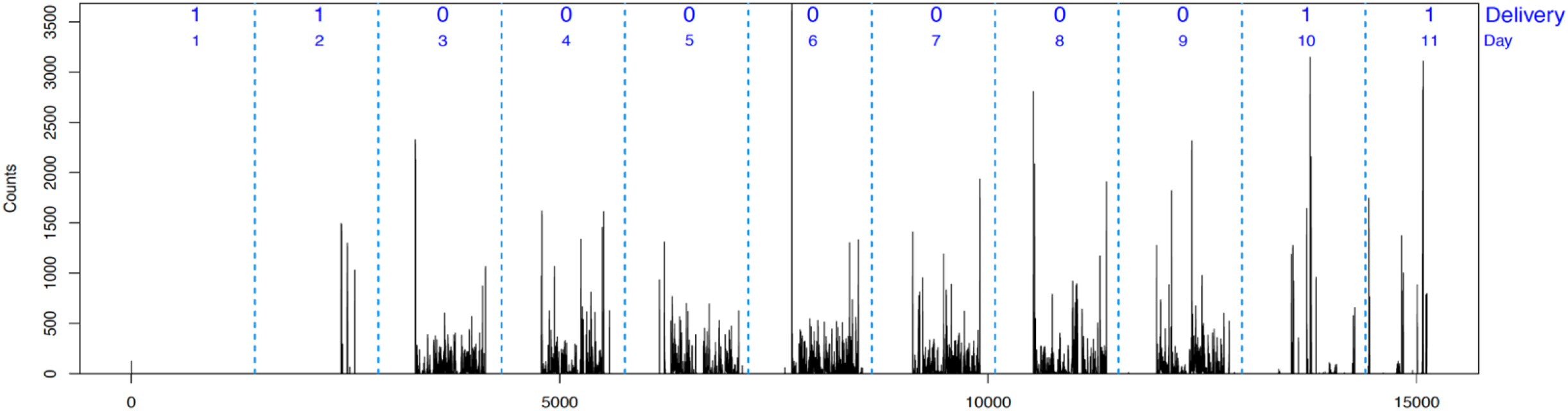
Example of accelerometry data for an assessment. The black lines represent the measurements on the x-axis wit h a one-minute epoch. Vertical dashed blue lines indicate midnight. The ‘delivery’ label indicates 0 for a human wear day and 1 for a delivery day. The blue text enumerates the day of the assessment.

The participants were requested to wear the Actigraph for the entire duration of the assessment, except when sleeping. Additionally, participants were requested to keep a timestamped log of when they received the Actigraph in the mail, when they returned the Actigraph to postal services, and any other potential issues such as non-adherence to protocol. These logs were used to label the days in the dataset as either human wear or delivery.

### Data Processing

Prior to modeling, we applied two methods to process the data. As the data were measured with 1-minute epoch, a complete day consists of 1440 measurements for each of the three axes. Every assessment contained at least one incomplete day during which less than 1440 measurements were taken. This occurred due to the accelerometer being activated or deactivated at any time other than midnight. Since the convolutional layers of a neural network require all inputs to be the same shape, the days that contained less than 1440 measurements were zero-padded (i.e., zero count at each minute) to a length of 1440. If the truncated day occurred at the start of the assessment, the zero-padding occurred from midnight to the time the Actigraph was activated. If the truncated day occurred at the end of the assessment, the zero-padding occurred from the deactivation time to the next midnight. Data that were only zero-padded were denoted as “minimally processed” data.

The data were also processed using procedures designed to remove days that contain little information. Any day was removed from the dataset if it had a total of less than 5000 counts or less than 10 minutes of movement in the vector magnitude of all three axes, which is calculated as the square root of the sum of squares of the count measurements from x, y, and z axes. Additionally, any day that was labeled as human wear was removed if less than 120 minutes of total activity occurred as this indicated a large amount of non-compliance with the protocol. The rational for this criteria is that days meeting these criteria would likely not be included in a typical data analysis as they do not meet criteria as qualified data (e.g., many studies require 600 minutes of wearing to be qualified as valid day). Data that were both zero-padded and processed with aforementioned criteria were denoted as “fully processed.” The difference in processing between the two methods is summarized in **Table 1**.

**Table 1:** Summary of methods to generate minimally and fully processed data.

| **Minimally Processed** | **Fully Processed** |
| --- | --- |
| Zeropad days with < 1440 minutes | Zeropad days with < 1440 minutes |
|  | Remove days with < 5000 total counts in vector magnitude |
|  | Remove days with < 120 minutes human activity |
|  | Remove days with < 10 minutes of delivery activity |

In this analysis, both the minimally and fully processed datasets were modeled in order to explore different algorithm’s capabilities of handling messier data. The minimally processed data approximates accelerometer data that is confounded with participant non-adherence, while the fully processed data is a much cleaner dataset. An example and visualization of the differences between the minimally and fully processed data is presented in **Figure 2**.

**Figure 2:**
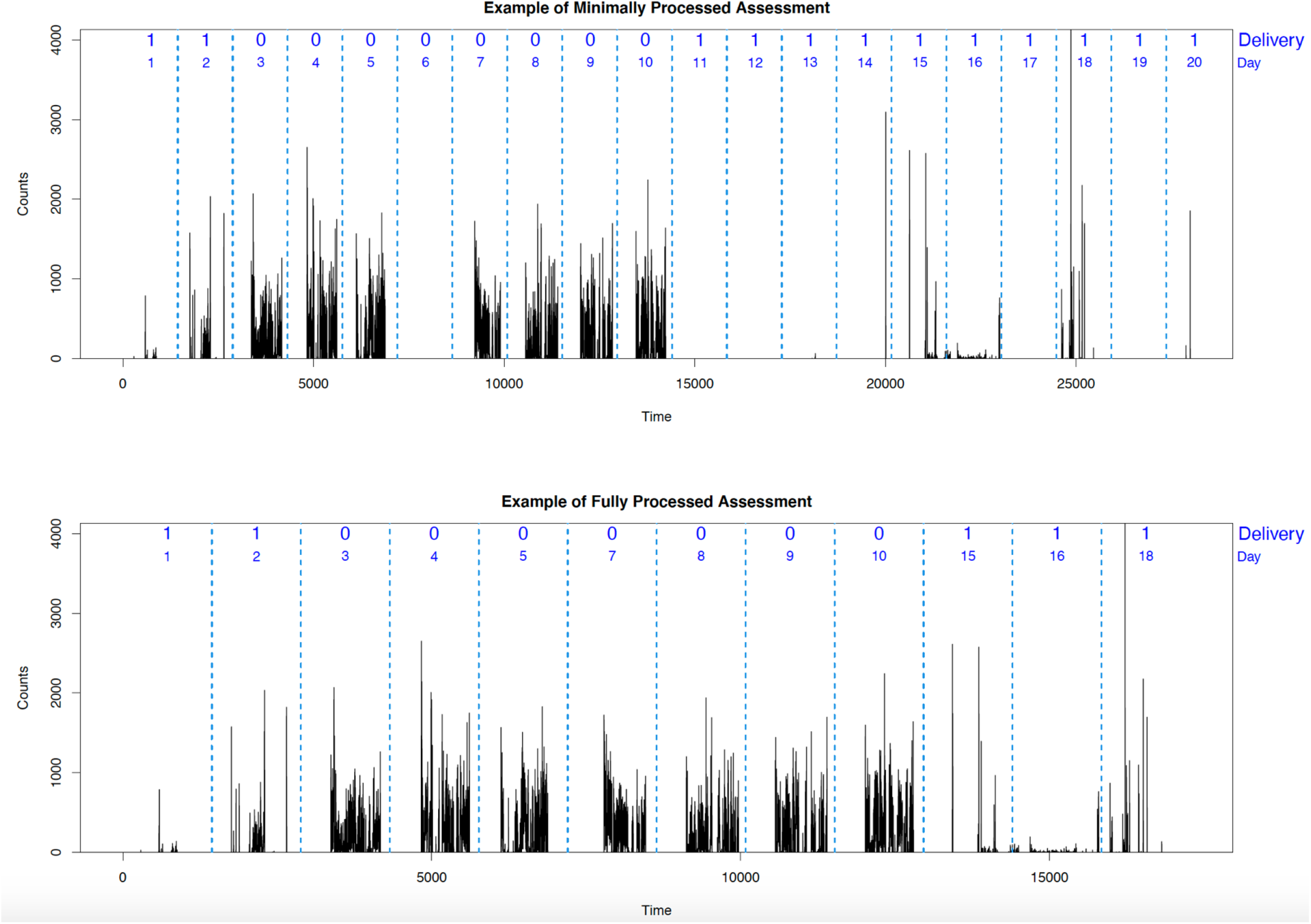
Example of minimally and fully processed data. No days are removed for the minimally processed data, while the non-compliance human activity day 6 is removed in the fully processed data. Additionally, the large stretch of several zero activity delivery days from day 11 to 14, as well as days 17 and 20 are removed in the full processing.

### Data Preparation and Feature Extraction

The data from each day were segmented into lengths of 1440 measurements between the hours of 0:00 and 23:59. These segments were reshaped into three dimensional arrays of stacks of 1440 by 3. The day long segments were used as inputs in the convolutional and recurrent neural networks or to extract features. We extracted 8 features from the vector magnitude, which include: mean, variance, maximum, 95th quantile, absolute energy, absolute change in energy, kurtosis, and skewness. The definition of absolute energy [1] and absolute change in energy [2] can be found below.

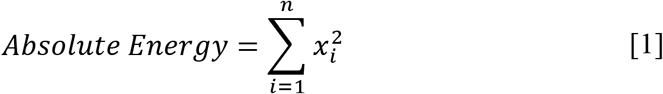

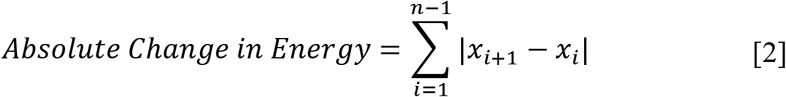

All the features and the raw data from the x, y, and z axes were mean centered and their standard deviation scaled, which is critical for achieving convergence in many of the models used in this analysis.

### Model Development

Seven different models were developed: random forest (Breiman, 2001), multi-layer perceptron (Rosenblatt, 1958), logistic regression, mixed-effects logistic regression, convolutional neural network (LeCun et al., 1999), LSTM recurrent neural network (Hochreiter and Schmidhuber, 1997), and a convolutional LSTM recurrent neural network (Shi et al., 2015). The random forest, regression models, and multi-layer perceptron utilized a traditional approach with extracted features from each data segment as inputs. On the other hand, the convolutional and recurrent neural networks used the scaled raw data as inputs in order to allow the models to learn local level features, but without including the extracted features. All neural networks were fit with a binary cross-entropy loss function and an Adam optimizer (Kingma and Ba, 2014) over 10 epochs of training.

The random forest model was developed using the 8 extracted features and was composed of 500 trees. Gini impurity was utilized as the criterion for measuring the quality of each split in the individual trees and the number of features considered at each split was the rounded down log base 2 of the number of total features (i.e., log_2_8 = 3) (Breiman, 2001).

The logistic and mixed-effects logistic regression models were also fit with the 8 extracted features, each of which was flexibly modeled with a restricted cubic spline with three knots. The mixed effects model was fit with a random intercept for participant.

The multi-layer perceptron (MLP) consists of 3 layers. The input layer is composed of 9 neurons - one for each of the 9 features. The input layer feeds into the first hidden layer, a dense layer consisting of 200 neurons with ReLU activation functions. This dense layer has 50% dropout as it feeds into the output layer, which is composed of a single output neuron with a sigmoid activation function. The architecture of the MLP is summarized in **Appendix, Table A1**.

The 1-D convolutional neural network consists of an input layer, output layer, and six hidden layers with ReLU activation functions. The input layer contains a single neuron for every measurement in a given day. The first hidden layer is a 1-D convolutional layer with 200 filters that span 5 measurements with a stride of 1. After a max pooling layer with a patch size of 4, the model has another pair of convolutional and max pooling layers that are composed of 64 filters with a span of 5 and a pooling size of 4, respectively. The output is then flattened and fed into a 64-neuron dense layer. The dense layer feeds into an output layer with a single sigmoid activated neuron. Details on the architecture can be found in **Appendix, Table A2**.

The recurrent neural network consists of an input layer, an output layer, and two hidden layers. The first hidden layer is composed of 30 long short-term memory cells with hyperbolic tangent activation functions and sigmoid recurrent activation functions. After a 40% dropout, the first hidden layer feeds into a dense layer of 100 neurons with ReLU activation functions. After 30% dropout, this layer fed into the output layer, which was composed of a single neuron with a sigmoid activation function. Details on the architecture can be found in **Appendix, Table A3**.

The convolutional recurrent neural network consists of an input layer, an output layer, and 4 hidden layers. The first and second hidden layers contain a 1-D convolutional layer and a max pooling layer with 200 filters of size 5 by 3 and a pooling size of 4 by 1, respectively. The 3rd hidden layer is another 1-D convolutional layer with 64 filters with a span of 5. Both convolutional layers had strides of 1 and ReLU activation functions. The 4th hidden layer is an LSTM layer with 30 memory cells with tanH activation functions and sigmoid recurrent activation functions. After 30% dropout, the LSTM layer fed into the output layer, which was composed of a single neuron with a sigmoid activation function. Details on the convolutional recurrent model architecture can be found in **Appendix, Table A4**.

### Model Assessment

Five-fold Monte Carlo cross-validation was performed to assess model performance. For each repetition, the test and training sets were selected by randomly sampling 30% and 70% of participants’ data, respectively. The data were split such that the training and test sets contained unique participants. Additionally, the centering and scaling of the raw data as well as feature extraction were performed separately for the test and training sets. The models were fit with the training sets, then the mean sensitivity, positive predictive value, F1 score, and Brier score (Brier, 1950) were calculated from the predictions of the test sets. Sensitivity [3] (also known as recall) is calculated as the ratio of true positive delivery classifications to true positives plus false negatives. Positive predictive value [4] (PPV) (also known as precision) is calculated as the ratio of true positives to true positives plus false positives. F1 score [5] is a commonly used general measure of model performance in the field of machine learning and is calculated as the harmonic mean of sensitivity and PPV. Brier score [6] is the mean square error of a model’s predicted probability, where *f_i_* indicates a model’s forecast and *o_i_* indicates the true outcome for *i^th^* sample across *N* samples.

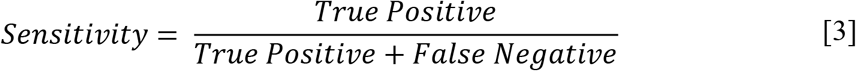

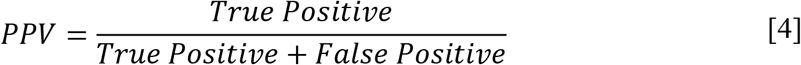

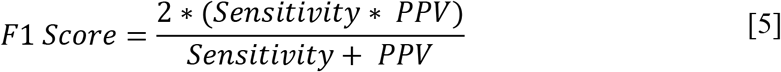

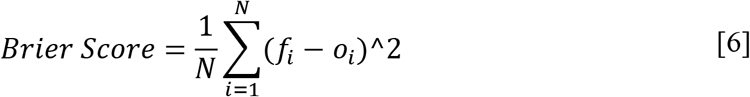

## Results

### Data Description

After minimal processing, the data had a total of 10,546 days, while the fully processed data had a total of 7,433 days. The days removed during human wear were likely caused by non-adherence. **Figure 3** shows that only 46% of the days in the minimally processed dataset are human wear, while 60% of the days in the fully processed dataset are human wear. Most of the days removed from full processing are delivery days with little or no activity.

**Figure 3:**
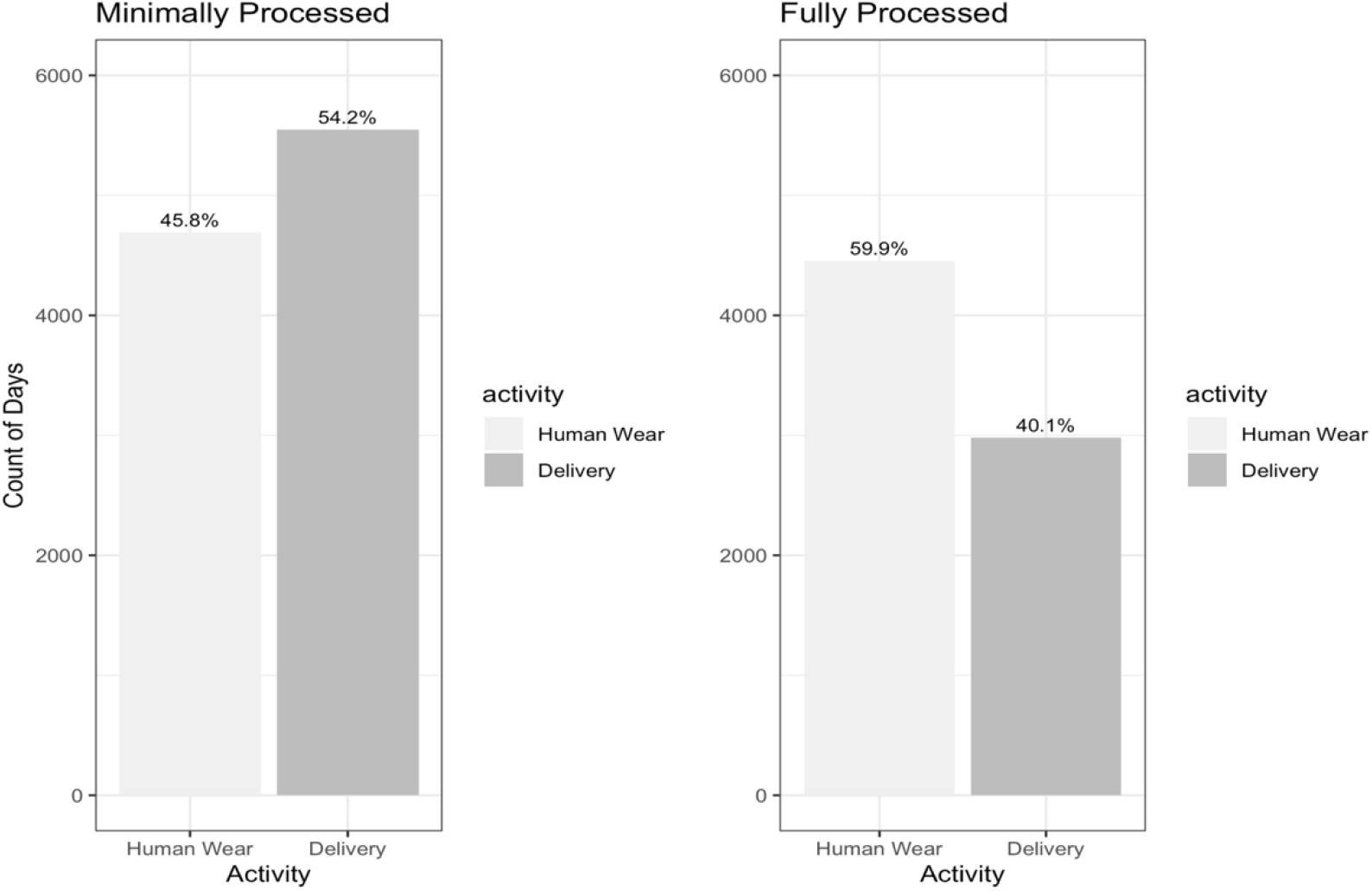
The number of days by activity in the minimally and fully processed datasets. The percentage of each activity is presented on top of each bar. The proportion of delivery days is decreased in the fully processed data.

Each subject participated in a range of one to three assessments in which they were asked to wear the Actigraph for a week. On average, the accelerometer for each assessment was active for approximately 17 days, much more than 7 days, suggesting many days are non-wear or delivery days. Across all assessments, the average number of days per participant was approximately 42 days. After fully processing the data, the average number of days per participant was reduced to approximately 30 days. The number of days by participant in the minimally and fully processed dataset is shown in **Figure 4**.

**Figure 4:**
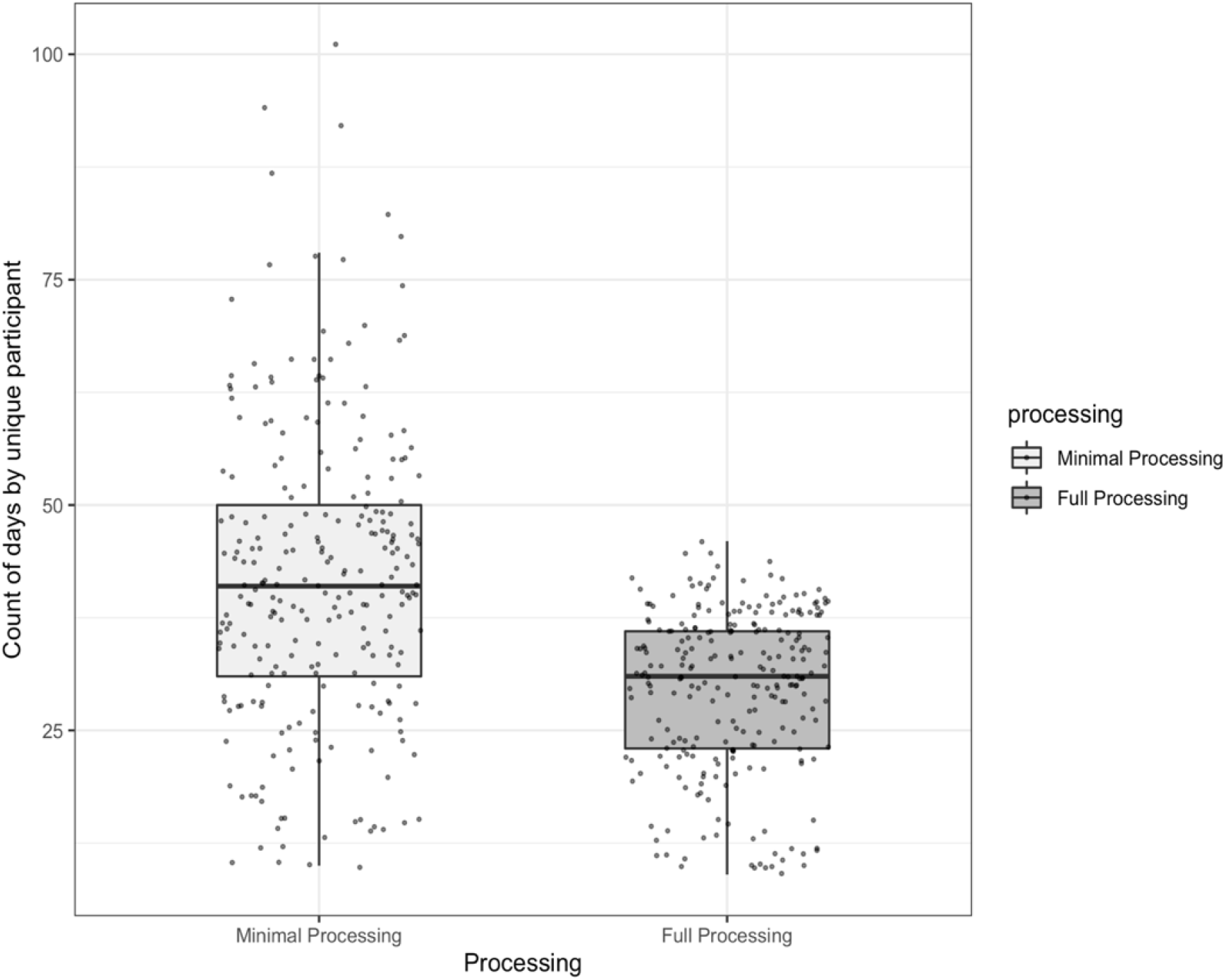
Box plot of the number of days by participant for the minimally and fully processed datasets. The center line of the boxplot indicates the median. The bottom and top hinges of the box indicate the 25^th^ and 75^th^ quantiles. The whiskers extend from the end of the box to a length of 1.5 multiplied by the interquartile range. Additionally, data points are overlaid on the boxplot.

### Model Performance

The mean of the sensitivity, PPV, F1 score, and Brier score across the 5 Monte Carlo cross-validations are presented in **Figure 5 (A)** and **(B)** for the minimally and fully processed data, respectively. The corresponding numerical results with both mean and standard deviation of the model performance metrics are also presented in a table in **Appendix, Table A5**. All models had a lower Brier score for the fully processed dataset compared to the minimally processed dataset; however, several models have a better F1 score in the minimally processed dataset. For both forms of processing, the recurrent architecture had the worst performance, while the convolutional recurrent neural network model marginally outperformed the other models with a mean F1 score of 0.961 and 0.960 in the minimally and fully processed datasets, respectively.

**Figure 5:**
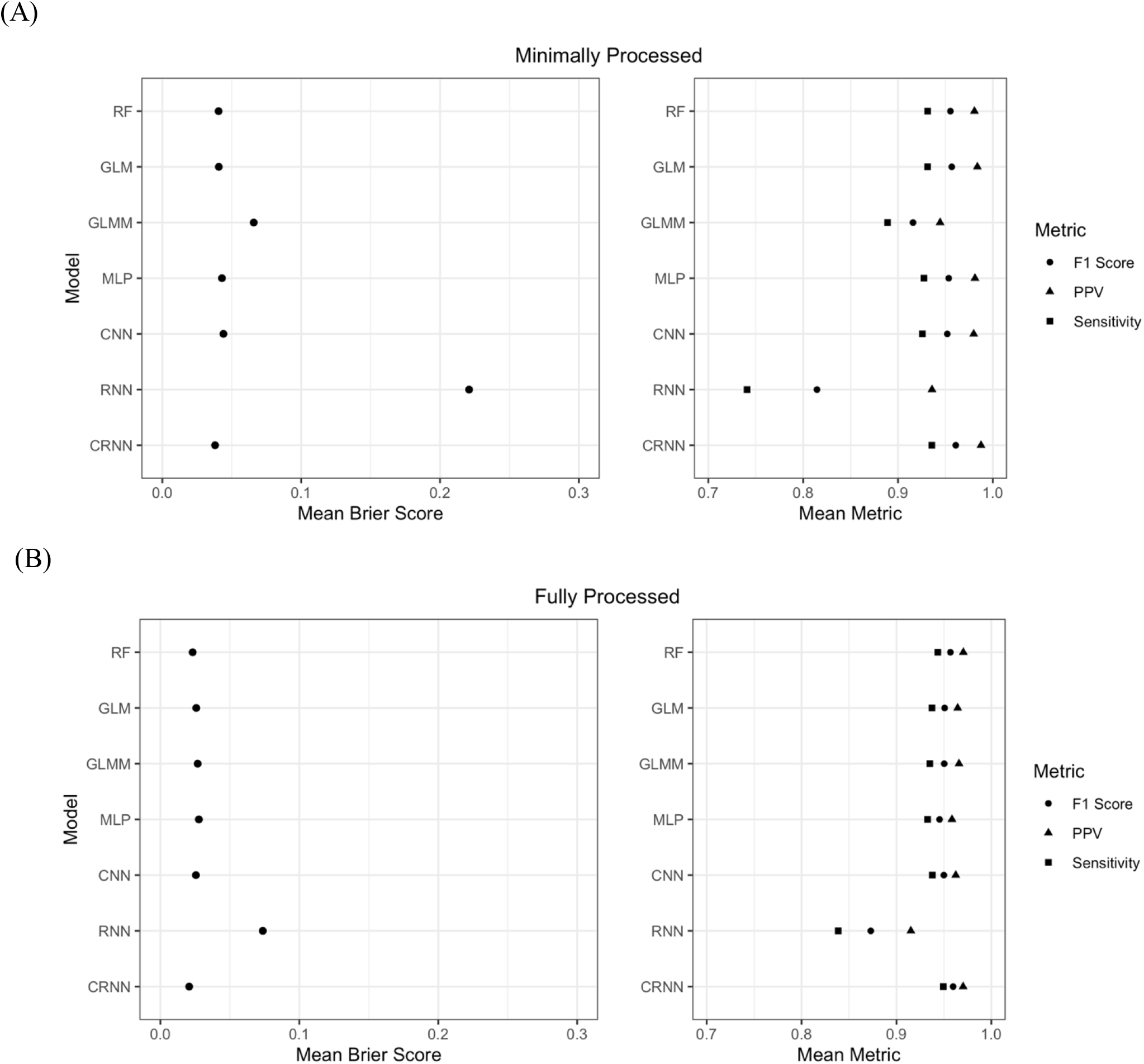
Cross-validated average model performance metrics on the minimally processed data (A), and fully processed data (B). RF: Random Forest; GLM: Generalized Linear Model; GLMM: Generalized Linear Mixed-Effects Model; MLP: Multilayer Perceptron; CNN: Convolutional Neural Network; RNN: Recurrent Neural Network; CRNN: Convolutional Recurrent Neural Network; PPV: Positive Predictive Value.

Out of the feature input models, the mixed-effects logistic regressions generally performed the worst, while the random forest marginally outperformed the other models in most of the metrics. The mixed-effect model performed fairly well when used on the fully processed dataset, but performed poorly relative to the other feature input models when used to model the minimally processed data. Out of the scaled raw data input models, the recurrent neural network performed the worst. Similarly to the mixed-effect model, the recurrent neural network’s performance was particularly poor relative to other models when used to model the minimally processed dataset. The convolutional neural network performed very well in both the minimally and fully processed data, but it was marginally outperformed by the convolutional recurrent neural network that performed best out of all the models.

## Discussion

This study used a large dataset of 10,546 days of activity with minimal processing and 7,433 days of activity after fully processing to develop models for the classification of days in accelerometry data as either human wear or delivery activity. We trained several statistical and machine learning models in a supervised learning context to discriminate between human wear and delivery days. All models performed well, especially with the fully processed data. A hybrid convolutional recurrent neural network marginally outperformed the other models with a mean 5-fold cross-validated Brier score of 0.021 and F1 score of 0.960. The logistic regression and random forest models also performed well with mean Brier scores of 0.026 and 0.023, and F1 scores of 0.951 and 0.957, respectively.

The convolutional and convolutional recurrent neural networks performed very well, while the recurrent neural network performed the worst out of all models using both the minimally processed and fully processed datasets. The slightly stronger performance of the convolutional recurrent neural network relative to the convolutional neural network indicates that incorporating time dependencies is helpful. However, the poor performance of the recurrent neural network indicates that the data greatly benefits from being reduced in dimensionality through convolutional layers before the recurrent layer processes the sequence. It is likely that the LSTM recurrent neural network has difficulties processing the thousands of days input with a length of 1440 measurements per day.

The convolutional recurrent neural network had the best performance, but the structure of the dataset makes the model somewhat naive in that it cannot differentiate between unique assessments. Ideally, the data would have been zero padded between assessments in order to reset the internal memory of the recurrent layers. Another potential improvement to the convolutional recurrent neural network would be the inclusion of a bidirectional LSTM layer. These layers incorporate information from both future and past states in an input sequence and have recently been shown to have improved performance over traditional LSTMs in certain contexts (Chiu & Nichols 2016).

The random forest and logistic regression marginally outperformed the multilayer perceptron model. The mixed effects model performed approximately as well as the random forest and logistic regression model for the fully processed data, but performed poorly when modeling the minimally processed data.

The largest limitation of this study is that the models were developed in a supervised learning context, and may not perform well with accelerometry data obtained from different studies. However, we expect reasonably good performance when the models are applied to new data considering the mechanistic nature of delivery activity and the high performance of our models during internal validation. Future work is need to externally validate the models with independent datasets.

Another important point to consider is an ease of each model’s implementation. The random forest and logistic regression models would be fairly simple to implement on a different dataset but do require certain statistical features to be extracted. One advantage of the feature extraction and scaling is that the models may be easily applicable to data with other temporal resolutions. The logistic regression model would be especially easy to import for use in any programming language as it has a closed form solution and would not require any package dependencies. Although the convolutional recurrent neural network showed the best performance for our dataset, it has a few barriers to widespread implementation. First, the model was trained on a dataset with a temporal resolution of one measurement per minute. The trained neural network would likely perform well for accelerometry datasets with the same temporal resolution but may not be generalizable to other resolutions since the model learned to recognize local features only at one-minute epoch. However, if the data was recorded at a higher resolution, it could easily be collapsed to a lower resolution using a program such as an R package, *PhysicalActivity* (Choi et al., 2018). Another issue in the widespread implementation of the convolutional recurrent neural network is its higher computational cost and dependency on the R package, *keras* (Chollet, 2015). Although it is fairly simple to export and import models, the requirement of both installing keras and running the model could deter some users.

## Conclusions

We developed several statistical and machine learning models that classify days in an accelerometry dataset as human wear or delivery activity with a high level of predictive accuracy. The majority of these models demonstrated excellent performance with both minimally and fully processed datasets. The top performing three models (random forest, logistic regression, and convolutional recurrent neural networks) and the processing techniques from this study are implemented in the R package, *PhysicalActivity*. Readers can find the most recent version of the R Package at https://github.com/couthcommander/PhysicalActivity. These models will allow an analyst to automate the cleaning of human activity accelerometry data that is adulterated with delivery data. In choosing the best model for application in identifying delivery days, the user can choose a model based on whether they want to use raw data or utilize manual feature extraction. The user can also weigh the higher computational cost and greater performance of the convolutional recurrent neural networks against the much faster but slightly less powerful random forest or logistic regression models. Future work is needed to externally validate the models with other datasets collected in diverse studies, and we welcome contributions from users.

## Acknowledgments

We thank Cole Beck for his assistance in making the program available in the R package, *PhysicalActivity*.

## Author Contributions

RM analyzed data and drafted the manuscript. LC conceived the research and drafted the manuscript. KRA provided the data and drafted the manuscript. All authors reviewed and edited the final manuscript.

## Appendix

**Table A1:**
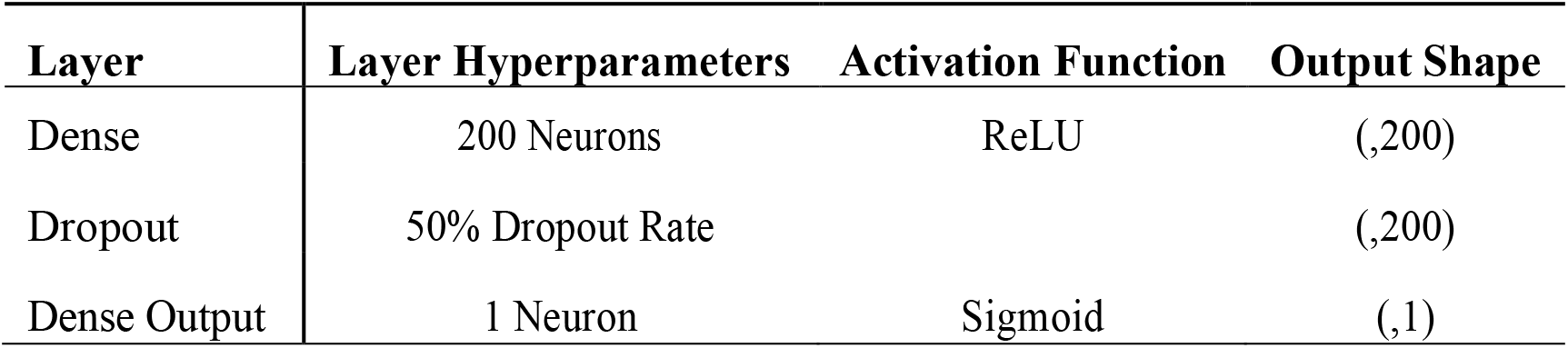
Architecture of multi-layer perceptron neural network.

**Table A2:**
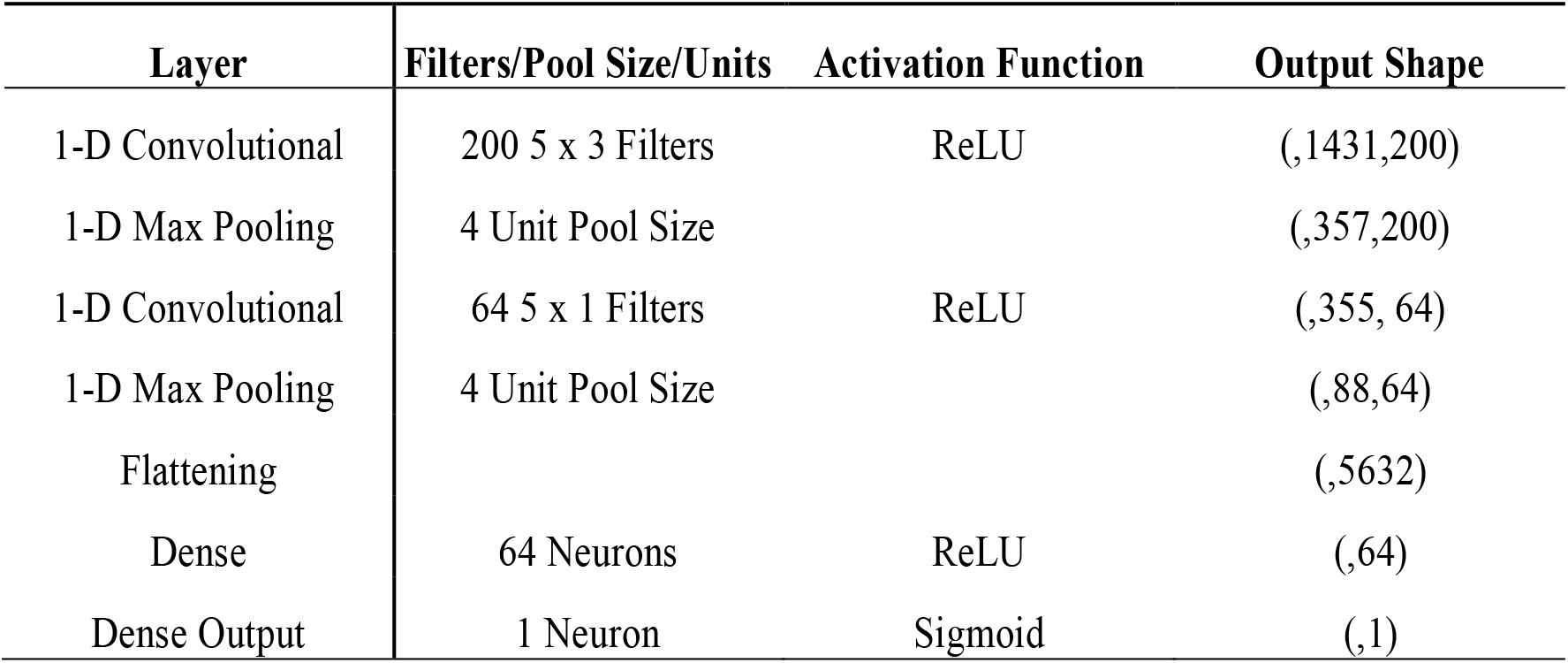
Architecture of 1-D convolutional neural network.

| Layer | Filters/Pool Size/Units | Activation Function | Output Shape |
| --- | --- | --- | --- |
| 1-D Convolutional | 200 5 x 3 Filters | ReLU | (,1431,200) |
| 1-D Max Pooling | 4 Unit Pool Size |  | (,357,200) |
| 1-D Convolutional | 64 5 x 1 Filters | ReLU | (,355, 64) |
| 1-D Max Pooling | 4 Unit Pool Size |  | (,88,64) |
| Flattening |  |  | (,5632) |
| Dense | 64 Neurons | ReLU | (,64) |
| Dense Output | 1 Neuron | Sigmoid | (,1) |

**Table A3:** Architecture of long short-term memory (LSTM) recurrent neural network.

| Layer | Layer Hyperparameters | Activation Function | Output Shape |
| --- | --- | --- | --- |
| LSTM | 30 Neurons | TanH and Sigmoid | (,30) |
| Dropout | 40% Dropout Rate |  | (,30) |
| Dense | 100 Neurons | ReLU | (,100) |
| Dropout | 30% Dropout Rate |  | (,100) |
| Dense Output | 1 Neuron | Sigmoid | (,1) |

**Table A4:** Architecture of convolutional long short-term memory (LSTM) neural network.

| Layer | Layer Hyperparameters | Activation Function | Output Shape |
| --- | --- | --- | --- |
| Convolutional | 200 5 x 3 Filters | ReLU | (,1431,200) |
| Max Pooling | 4 Unit Pool Size |  | (,357,200) |
| Convolutional | 64 5 x 1 Filters | ReLU | (,355,64) |
| LSTM | 30 Neurons | TanH and Sigmoid | (,30) |
| Dropout | 30% Dropout Rate |  | (,30) |
| Dense Output | 1 Neuron | Sigmoid | (,1) |

**Table A5:**
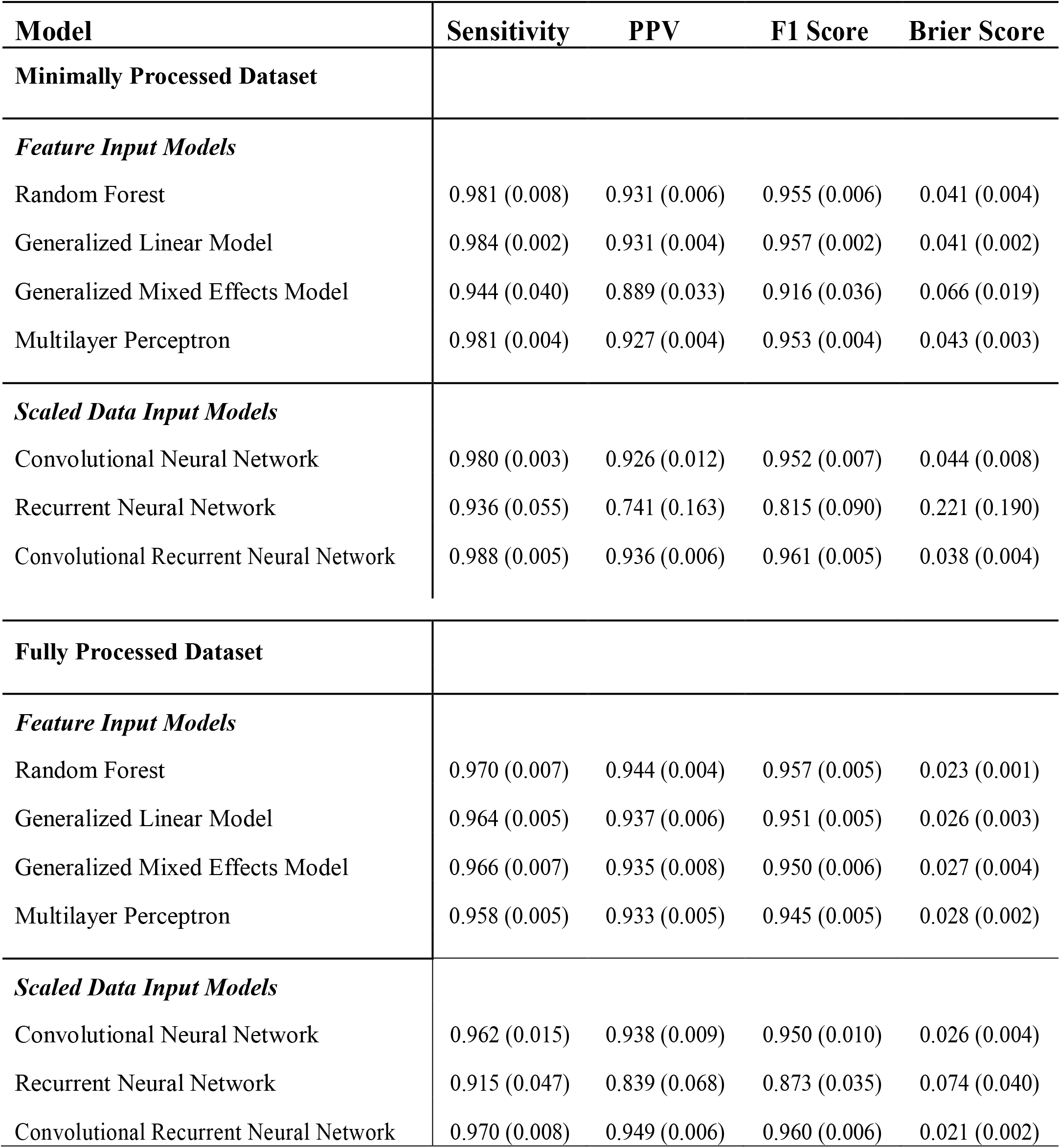
Average model performance metrics from 5-fold Monte Carlo cross-validation with standard deviation in parentheses for the minimally and fully processed data. PPV: Positive Predictive Value.

| **Model** | **Sensitivity** | **PPV** | **F1 Score** | **Brier Score** |
| --- | --- | --- | --- | --- |
| **Minimally Processed Dataset** |  |  |  |  |
| ***Feature Input Models*** |  |  |  |  |
| Random Forest | 0.981 (0.008) | 0.931 (0.006) | 0.955 (0.006) | 0.041 (0.004) |
| Generalized Linear Model | 0.984 (0.002) | 0.931 (0.004) | 0.957 (0.002) | 0.041 (0.002) |
| Generalized Mixed Effects Model | 0.944 (0.040) | 0.889 (0.033) | 0.916 (0.036) | 0.066 (0.019) |
| Multilayer Perceptron | 0.981 (0.004) | 0.927 (0.004) | 0.953 (0.004) | 0.043 (0.003) |
| ***Scaled Data Input Models*** |  |  |  |  |
| Convolutional Neural Network | 0.980 (0.003) | 0.926 (0.012) | 0.952 (0.007) | 0.044 (0.008) |
| Recurrent Neural Network | 0.936 (0.055) | 0.741 (0.163) | 0.815 (0.090) | 0.221 (0.190) |
| Convolutional Recurrent Neural Network | 0.988 (0.005) | 0.936 (0.006) | 0.961 (0.005) | 0.038 (0.004) |
| **Fully Processed Dataset** |  |  |  |  |
| ***Feature Input Models*** |  |  |  |  |
| Random Forest | 0.970 (0.007) | 0.944 (0.004) | 0.957 (0.005) | 0.023 (0.001) |
| Generalized Linear Model | 0.964 (0.005) | 0.937 (0.006) | 0.951 (0.005) | 0.026 (0.003) |
| Generalized Mixed Effects Model | 0.966 (0.007) | 0.935 (0.008) | 0.950 (0.006) | 0.027 (0.004) |
| Multilayer Perceptron | 0.958 (0.005) | 0.933 (0.005) | 0.945 (0.005) | 0.028 (0.002) |
| ***Scaled Data Input Models*** |  |  |  |  |
| Convolutional Neural Network | 0.962 (0.015) | 0.938 (0.009) | 0.950 (0.010) | 0.026 (0.004) |
| Recurrent Neural Network | 0.915 (0.047) | 0.839 (0.068) | 0.873 (0.035) | 0.074 (0.040) |
| Convolutional Recurrent Neural Network | 0.970 (0.008) | 0.949 (0.006) | 0.960 (0.006) | 0.021 (0.002) |

## References

Bagalà F, Becker C, Cappello A, Chiari L, Aminian K, Hausdorff JM, et al. Evaluation of accelerometer-based fall detection algorithms on real-world falls. PLoS ONE. 2012;7(5):e37062.

Crouter SE, Churilla JR, Bassett DR. Estimating energy expenditure using accelerometers. Eur J Appl Physiol. 2006 Dec;98(6):601–12.

Kwapisz JR, Weiss GM, Moore SA. Activity recognition using cell phone accelerometers. SIGKDD Explor Newsl. 2011 Mar 31;12(2):74–82.

Choi L, Liu Z, Matthews CE, Buchowski MS. Validation of Accelerometer Wear and Nonwear Time Classification Algorithm. Med Sci Sports Exerc. 2011 Feb;43(2):357–364.

Choi L, Ward SC, Schnelle JF, Buchowski MS. Assessment of Wear/Nonwear Time Classification Algorithms for Triaxial Accelerometer. Med Sci Sports Exerc. 2012 Oct;44(10):2009–16.

Kim E, Helal S, Cook D. Human Activity Recognition and Pattern Discovery. IEEE Pervasive Comput. 2010;9(1):48.

Lara OD, Labrador MA. A Survey on Human Activity Recognition using Wearable Sensors. IEEE Communications Surveys Tutorials. 2013 Third;15(3):1192–209.

Rosenblatt F. The perceptron: A probabilistic model for information storage and organization in the brain. Psychological Review. 1958;65(6):386–408.

Ellis K, Kerr J, Godbole S, Lanckriet G, Wing D, Marshall S. A random forest classifier for the prediction of energy expenditure and type of physical activity from wrist and hip accelerometers. Physiol Meas. 2014 Nov;35(11):2191–203.

Kingma D, Ba J. Adam: A method for stochastic optimization. arXiv. 2014 Dec 22;1412:6980.

Breiman L. Random Forests. Machine Learning. 2001 Oct 1;45(1):5–32.

LeCun Y, Haffner P, Bottou L, Bengio Y. Object Recognition with Gradient-Based Learning. In: Forsyth DA, Mundy JL, di Gesú V, Cipolla R, editors. Shape, Contour and Grouping in Computer Vision [Internet]. Berlin, Heidelberg: Springer; 1999 [cited 2020 Jul 24]. p. 319–45. (Lecture Notes in Computer Science). Available from:https://doi.org/10.1007/3-540-46805-6_19

Hochreiter S, Schmidhuber J. Long Short-Term Memory. Neural Computation. 1997 Nov 1;9(8):1735–80.

Ignatov A. Real-time human activity recognition from accelerometer data using Convolutional Neural Networks. Applied Soft Computing. 2018 Jan 1;62:915–22.

Ordóñez F, Roggen D. Deep Convolutional and LSTM Recurrent Neural Networks for Multimodal Wearable Activity Recognition. Sensors. 2016 Jan 18;16(1):115.

Archer KR, Haug CM, Pennings J. Combining Two Programs to Improve Disability, Pain, and Health Among Patients Who Have Had Back Surgery. Washington, DC: Patient-Centered Outcomes Research Institute (PCORI). 2020

Coronado RA, Robinette PE, Henry AL, Pennings JS, Haug CM, Skolasky RL, et al. Bouncing back after lumbar spine surgery: early postoperative resilience is associated with 12-month physical function, pain interference, social participation, and disability. The Spine Journal. 2020 Jul;S1529943020310020.

Shi X, Chen Z, Wang H, Yeung D-Y, Wong W, Woo W. Convolutional LSTM Network: A Machine Learning Approach for Precipitation Nowcasting. In: Cortes C, Lawrence ND, Lee DD, Sugiyama M, Garnett R, editors. Advances in Neural Information Processing Systems 28[Internet]. Curran Associates, Inc.; 2015 [cited 2020 Aug 11]. p. 802–810. Available from: http://papers.nips.cc/paper/5955-convolutional-lstm-network-a-machine-learning-approach-for-precipitation-nowcasting.pdf

Brier GW. Verification of Forecasts Expressed in Terms of Probability. Monthly Weather Review. 1950 Jan 1;78(1):1–3.

Chiu JPC, Nichols E. Named Entity Recognition with Bidirectional LSTM-CNNs. arXiv:151108308 [cs] [Internet]. 2016 Jul 19 [cited 2020 Jul 22]; Available from: http://arxiv.org/abs/1511.08308

Choi L, Beck C, Liu Z, Matthews CE and Buchowski MS. PhysicalActivity Process Accelerometer Data for Physical Activity Measurement. R package version 0.2-2. 2018; Available from:https://CRAN.R-project.org/package=PhysicalActivity

Chollet, F. Keras. Github Repository. 2018; Available from:https://github.com/fchollet/keras

